# Polygenic Risk Score of Sporadic late Onset Alzheimer Disease Reveals a Shared Architecture with the Familial and Early Onset Forms

**DOI:** 10.1101/165209

**Authors:** Jorge L Del-Aguila, Benjamin Saef, Kathleen Black, Maria Victoria Fernandez, John Budde, Laura Ibanez, Manav Kapoor, Giuseppe Tosto, Richard P Mayeux, David M Holtzman, Anne M. Fagan, John C Morris, Randall J. Bateman, Alison Goate, the Dominantly Inherited Alzheimer Network (DIAN), Disease Neuroimaging Initiative (ADNI), the NIA-LOAD family study, Carlos Cruchaga, Oscar Harari

**Author notes:** Joint First Authors who contributed equally (Jorge L Del-Aguila, Benjamin Saef). Authors with same contribution (Carlos Cruchaga, Oscar Harari). Data used in preparation of this article were obtained from the Alzheimer’s Disease Neuroimaging Initiative (ADNI) database (adni.loni.usc.edu). As such, the investigators within the ADNI contributed to the design and implementation of ADNI and/or provided data but did not participate in analysis or writing of this report. A complete listing of ADNI investigators can be found at: http://adni.loni.usc.edu/wpcontent/uploads/how_to_apply/ADNI_Acknowledgement_List.pdf. To whom correspondence should be addressed: Oscar Harari, PhD, Department of Psychiatry, Washington University School of Medicine, 660 South Euclid Avenue B8134, St. Louis, MO 63110.

## Abstract

**Objective:** To determine whether the genetic architecture of sporadic late-onset Alzheimer’s Disease (sLOAD) has an effect on familial late-onset AD (fLOAD), sporadic early-onset (sEOAD) and autosomal dominant early-onset (eADAD).

**Methods:** Polygenic risk scores (PRS) were constructed using previously identified 21 genome-wide significant loci for LOAD risk.

**Results:** We found that there is an overlap in the genetic architecture among sEOAD, fLOAD, and sLOAD. sEOAD showed the highest odds for the PRS (OR=2.27; p=1.29×10^-7^), followed by fLOAD (OR=1.75; p=1.12×10^-7^) and sLOAD (OR=1.40; p=1.21×10^-3^). PRS is associated with cerebrospinal fluid ptau_181_-Aβ_42_ on eADAD.

**Conclusion:** Our analysis confirms that the genetic factors identified for sLOAD also modulate risk in fLOAD and sEOAD cohorts. Furthermore, our results suggest that the burden of these risk variants is associated with familial clustering and earlier-onset of AD. Although these variants are not associated with risk in the eADAD, they may be modulating age at onset.

## Introduction

Alzheimer Disease (AD) is the most common form of dementia. In AD, the onset of cognitive impairment is preceded by a long preclinical phase, lasting approximately 15 to 20 years [1]. There is a large variability in the age at onset of AD, and only a small fraction of cases (1%) present clinical symptoms at an early age at onset (EO: before age 65). AD has a substantial but heterogeneous genetic component. Mutations in the *Amyloid-beta Precursor Protein* (*APP*) and *Presenilin* genes [2-6] cause the Mendelian forms of AD. Although autosomal dominant AD typically is associated with early symptoms onset (eADAD), some families that carry known pathogenic mutations present a late onset (onset>65-70) [7], suggesting a continuum between late and early onset. Additionally, a large proportion of AD cases with strong familial history of dementia also present a late-onset, and a complex genetic architecture (familial LOAD; fLOAD).

Most of the sporadic AD cases present a late onset (sLOAD), but occasionally can present an early onset (sEOAD). The *APOE* ε4 allele increases risk for sporadic early -and late-onset AD [6] and also for familial late onset AD [7, 8] (3 fold effect size for heterozygous carriers and 12 fold for homozygous carriers [9]). More recent genome-wide association studies of sporadic late onset AD have identified additional loci with moderate protective and risk effects [10-15]. Further studies suggest that Polygenic Risk Scores (PRS) created based on the 21 genome-wide loci capture the overall genetic architecture of late-onset AD and may help to predict AD risk [8, 16, 17].

PRS aggregates the effects that multiple genetic markers (both protective and risk variants) confer to individuals for a specific complex trait [18]. When employed as biomarkers, PRS can provide important insights about the prognosis of the disease, and can highlight early intervention strategies as well as inclusion criteria for targeted enrollment in clinical trials. Furthermore, PRS can be employed as a measure to identify the extent of overlap between the genetic architecture of co-morbid complex traits [19]. This is done, by evaluating the pleiotropic effects that the markers identified in one trait have in another trait, usually evaluated in an independent cohort. For example, this approach has been employed to study the shared genetic architecture between schizophrenia and cognitive function, as well as between depressive disorder and body mass index [19].

It is unknown whether the GWAS loci identified for sporadic LOAD have any effect on additional cases (sEOAD, fLOAD or eADAD), or the relative genetic burden of those variants in the other classifications of the disease. Therefore, a thorough evaluation of these variants will help us understand the extent of the genetic architecture shared among the different classifications of AD. This will also be informative for current preventative drug trials, some of which are focused on Mendelian forms or individuals with high genetic risk under the assumption that there are global pathways which are similar in early and late onset AD.

We tested the hypothesis that features of the genetic architecture identified for sLOAD are shared by participants with other clinical manifestations of AD. To do so, we derived the PRS from common variants identified in GWAS of *s*LOAD [15] in additional cohorts of affected participants with European ancestry with early onset AD (both sEOAD and eADAD) as well as fLOAD. Then we tested the association of the PRS with the clinical status in each of these cohorts. Finally, we explored whether the PRS is modulating additional aspects of AD, and evaluated its association with the age at onset.

## Materials and Methods

### Samples

We included participants with European ancestry from the Knight-Alzheimer’s Disease Research Center (Knight-ADRC) and the Dominantly Inherited Alzheimer Network (DIAN) study at Washington University [20], the Alzheimer’s Disease Neuroimaging Initiative (ADNI)[21], and the National Institute on Aging Genetics Initiative for Late-Onset Alzheimer’s Disease (NIA-LOAD)[22].

<u>Cohorts: eADAD</u> are defined as affected participants that carry known highly penetrant mutations in the *Presenilin* or APP genes. All the samples were selected from the DIAN study. <u>fLOAD:</u> Includes affected subjects from families with recorded familial history. Only one proband per family were included in the study. Probands were required to have a diagnosis of definite or probable late-onset AD (onset >65 years) and a sibling with definite, probable or possible late-onset AD with a similar age at onset. A third biologically-related family member (first, second or third degree) was also required, regardless of cognitive status. All the samples were selected from the NIA-LOAD study. <u>sEOAD</u> were defined as participants with diagnosis of AD, with an age at onset <65 without documented familial-history of AD. Samples were selected from the Knight-ADRC and ADNI. <u>sLOAD:</u> were defined as participants with clinical diagnosis of AD, age at onset >65, that do not specify documented family history of AD. Samples were selected from the Knight-ADRC and ADNI. <u>Controls</u> were defined as individuals older than 65 that after neurological assessment were determined to be non-affected. Unrelated samples were selected from the Knight-ADRC and ADNI. We included 236 sEOAD, 1021 sLOAD and 687 controls from the Knight-ADRC; 122 *s*EOAD, 226 sLOAD and 324 cognitively normal controls from ADNI; 1220 unrelated fLOAD from the NIA-LOAD and 249 eADAD (mutation carriers from the DIAN study).

<u>Description of the AD datasets:</u> Knight-ADRC research participants were evaluated by Clinical Core personnel at Washington University. Cases received a clinical diagnosis of AD in accordance with standard criteria and the presence or absence AD symptoms, and when present, their severity, was determined using the Clinical Dementia Rating (CDR)[23]. Neuropsychological and clinical assessments and biological samples were collected for all participants as described previously [24-29]. ADNI individuals were evaluated as described in the ADNI procedures manual (http://www.adni-info.org). Participants from the National Institute of Aging Late Onset Alzheimer Disease Family Study (NIA-LOAD Family Study), all cases had been diagnosed with AD dementia using criteria equivalent to the National Institute of Neurological and Communication Disorders and Stroke-Alzheimer’s Disease and Related Disorders Association (NINCDS-ADRDA) for probable AD [30]. All individuals had a family history of Alzheimer’s disease. Probands were required to have a diagnosis of definite or probable late-onset AD (onset >60 years) and a sibling with definite, probable or possible late-onset AD with a similar age at onset. A third biologically-related family member (first, second or third degree) was also required, regardless of cognitive status. If unaffected, this individual had to be ≥60 years of age, but ≥50 years of age if diagnosed with LOAD or mild cognitive impairment. Within each pedigree, we selected a single individual to screen by identifying the youngest affected family member with the most definitive diagnosis (i.e. individuals with autopsy confirmation were chosen over those with clinical diagnosis only). Finally, eADAD participants (n=249) were drawn from the Dominantly Inherited Alzheimer Network (DIAN)[1, 31]. Individuals at risk for carrying a mutation for autosomal dominant AD (*i.e. PSEN1, PSEN2* or *APP*) mutations] were enrolled in DIAN study. Participants from families with known pathogenic eADAD mutations were recruited from 197 families at six sites in the USA, one in the UK and three in Australia [1, 31]. The process of recruitment and enrolment has been described in detail previously [1, 31].

### Genotyping Platforms and Proxy selection

Table 1 summarizes the demographic and clinical characteristics of each clinical group. The Institutional Review Board of all participating institutions approved the study. Written informed consent was obtained from participants or their family members.

**Table 1.** Demographics of the cohorts

| Cohort | N | Female % | Age (SD)* | APOE ε4+(%)<sup>†</sup> |
| --- | --- | --- | --- | --- |
| Autosomal Dominant Early Onset (DIAN) | 249 | 57.03 | 37.16 (14.03) | 27.18 |
| Sporadic Early Onset<sub>3</sub> | 358 | 49.17 | 60.41 (4.83) | 62.74 |
| Familial Late Onset | 1220 | 63.10 | 75.30 (8.34) | 74.08 |
| Sporadic Late Onset | 1247 | 57.30 | 76.18(7.00) | 58.27 |
| Non-demented (Controls) | 1011 | 56.38 | 77.03 (7.05) | 31.22 |

\* Age at onset for affected participants and age of last assessment for non-demented subjects, and for DIAN participants age of recruitment
<sup>†</sup> Percentage of participants carriers of APOE ε4 allele.

The fLOAD cohort is part of NIA-LOAD and were genotyped as describe previously [32]. Participants, from the Knight-ADRC, DIAN and ADNI, were genotyped with the Illumina 610 chip, Omni-express chip or HumanCore Exome (Illumina, San Diego, CA, USA). All samples were imputed using SHAPEIT/IMPUTE2 [33, 34] with the 1000 Genomes Project as reference panel [35]. We discarded genotypes that did not pass quality criteria, and retained those with dosage levels >0.9 across all three genotype possibilities, and an information score filter >0.3.

In addition, we performed gender verification, tested for duplicates and unexpected familial relatedness, which were removed from our analysis, by estimating the pairwise genome-wide estimates of the proportion identity-by-descent. We performed standard quality control procedures on each genotyping array separately before combining data. A call rate ≥ 98% was applied for SNPs and individuals. SNPs not in Hardy-Weinberg equilibrium (p<1×10^−6^) or with MAF < 0.02 were excluded. We inferred the population structure and confirmed the ethnicity of participants by calculating the principal components using PLINK v1.9 (http://www.cog-genomics.org/plink2). Apolipoprotein E (*APOE*) genotype was determined for all individuals [28]. Briefly, *APOE* ε2, ε3, and ε4 isoforms were determined by genotyping rs7412 and rs429358 using Taqman genotyping technology as previously described [28, 36].

We employed proxy SNPs to tag the genome-wide significant loci reported for sLOAD [15] that did not pass our quality control process. We selected the SNPs with the highest genotyping rates, the highest R^2^ and D’ values (in both imputed data and in the 1000 genomes) to the reported SNPs in the IGAP study [15].

### Computation of Polygenic Risk Score

We derived a weighted PRS [18], modeling the odd ratios as reported in IGAP study [15] using a logarithm of base 2 transformation. SNPs utilized for the score would either need to have a high genotyping rate (greater than 90%) or otherwise be a reasonable proxy to the IGAP hits. We utilized PLINK v1.90b3.42 to calculate the PRS choosing the score function and the no-mean-imputation option to ensure that no scores would be imputed. The resulting mean was corrected by the multiplying allele count (Log OR Score).

### Statistical Analysis

The effect and statistical significance of the PRS were ascertained using logistic regression (The R Foundation for Statistical Computing v3.3.1) on the extreme tertiles (R quantile function). We have employed multiple models to investigate the extent of overlap of the genetic architecture of AD under different scenarios. The model 1 includes the calculated PRS correcting for sex, study as well as age for the late onset cohorts. The effects of *APOE* ε2 and ε4 genotypes are evaluated in the model 2, which extends the PRS to include the logarithm base 2 transformation of the odds ratios for APOE genotypes (i.e. ε2/ε2=0.6, ε2/ε3=0.6, ε2/ε4 =2.6, ε3/ε4=3.2, ε4/ε4=14.9) as previously reported [9]. Sex, study and age (for the late-onset cohorts) are also included as covariates in the model 2. To analyze the eADAD cohort we employed the functions glmer and lmer (package gee 4.13.19) clustering at family level to ascertain the effect of the PRS on clinical status and CSF biomarkers respectively. The statistical significance between the effects of the cohorts was calculated deriving the Z-score of the absolute difference of the odd ratios, corrected by its standard error.

The area under the curve (AUC) for receiver operating characteristic (ROC) analysis was calculated using the R package pROC v1.8, correcting for the same covariates as the logistic model. The areas obtained for the models were compared using the function roc.test.

Quantile regression [37] models for the relation between the PRS and the age at onset for sporadic early and LOAD participants were calculated for the 5-quantiles (quintiles) using the R package quantreg v5.29.

### Analyte Measurement

CSF tau, ptau_181_, and Aβ_42_ (markers of neuronal injury, neurofibrillary tangles and amyloid, respectively) were measured in 805 individuals enrolled in studies at the Knight-ADRC, 787 individuals from ADNI (390 from ADNI1 and 397 from ADNI2). Both studies measured biomarker values with internal standards and controls [38, 39]. However, there are differences between the measured values for each study due to differences in the antibodies and measurement technologies used by each center. CSF values were normalized as previously described [28, 40] before analyses.

### ADNI material and methods

Data used in the preparation of this article were obtained from the ADNI database (www.loni.ucla.edu\ADNI). The ADNI was launched in 2003 by the National Institute on Aging, the National Institute of Biomedical Imaging and Bioengineering, the Food and Drug Administration, private pharmaceutical companies and non-profit organizations, as a $60 million, 5-year public-private partnership. The Principal Investigator of this initiative is Michael W. Weiner, M.D. ADNI is the result of efforts of many co-investigators from a broad range of academic institutions and private corporations, and subjects have been recruited from over 50 sites across the U.S. and Canada. The initial goal of ADNI was to recruit 800 adults, ages 55 to 90, to participate in the research -approximately 200 cognitively normal older individuals to be followed for 3 years, 400 people with MCI to be followed for 3 years, and 200 people with early AD to be followed for 2 years.” For up-to-date information see www.adni-info.org.

## Results

### The PRS derived from genome-wide meta-analysis studies has similar effect for late-onset sporadic and familial AD

We initially verified the prediction accuracy of the PRS by ascertaining the cohort of *s*LOAD from the Knight-ADRC and ADNI participants. We observed that the PRS is significantly associated with clinical status for the sLOAD cases (model 1: OR=1.40; *p*=1.21×10^-3^; **Table 2**). As *APOE* alleles have a strong effect on the *s*LOAD, we extended the PRS to reflect the risk and protection effects of the ε4 and ε2 genotypes. The addition of the APOE genotype increased the effect of the PRS (model 2: OR=4.01; *p*=5.29×10^-34^). The ROC analysis revealed an Area Under the Curve (AUC= 0.67; 95% CI= 0.65-0.69) (**Figure 1**), which resembles the results of other studies [8, 17].

**Table 2.** Association results of the logistic regression models for Polygenic Risk Scores derived for each of the cohorts and compared to non-demented participants*

| Cohort | PRS<br>(model 1) |  | PRS w/APOE effects<br>(model 2) |  |
| --- | --- | --- | --- | --- |
|  | OR | *p-value* | OR | *p-value* |
| Autosomal Dominant Early Onset AD | 0.96 | $9.73 \times 10^{-1}$ | 0.98 | $9.66 \times 10^{-1}$ |
| Sporadic Early Onset AD | 2.27 | $1.29 \times 10^{-7}$ | 6.44 | $5.80 \times 10^{-26}$ |
| Familial Late Onset AD | 1.75 | $1.12 \times 10^{-7}$ | 7.85 | $2.52 \times 10^{-48}$ |
| Sporadic Late Onset AD | 1.40 | $1.21 \times 10^{-3}$ | 4.01 | $5.29 \times 10^{-34}$ |

\* The models correct for the sex and contributing study for the Early Onset cohorts and also age at onset or age of last assessment for the Late Onset cohorts. The PRS was analyzed by itself, as well as the effects of APOE genotypes in the cohorts. An extended version of the PRS adds the risk conferred by APOE into the effects. These results are a product of analyzing the tertiles with lowest and highest PRS values and a logistic regression for the entire set of samples in the case of models for APOE and no PRS.

**Figure 1.**
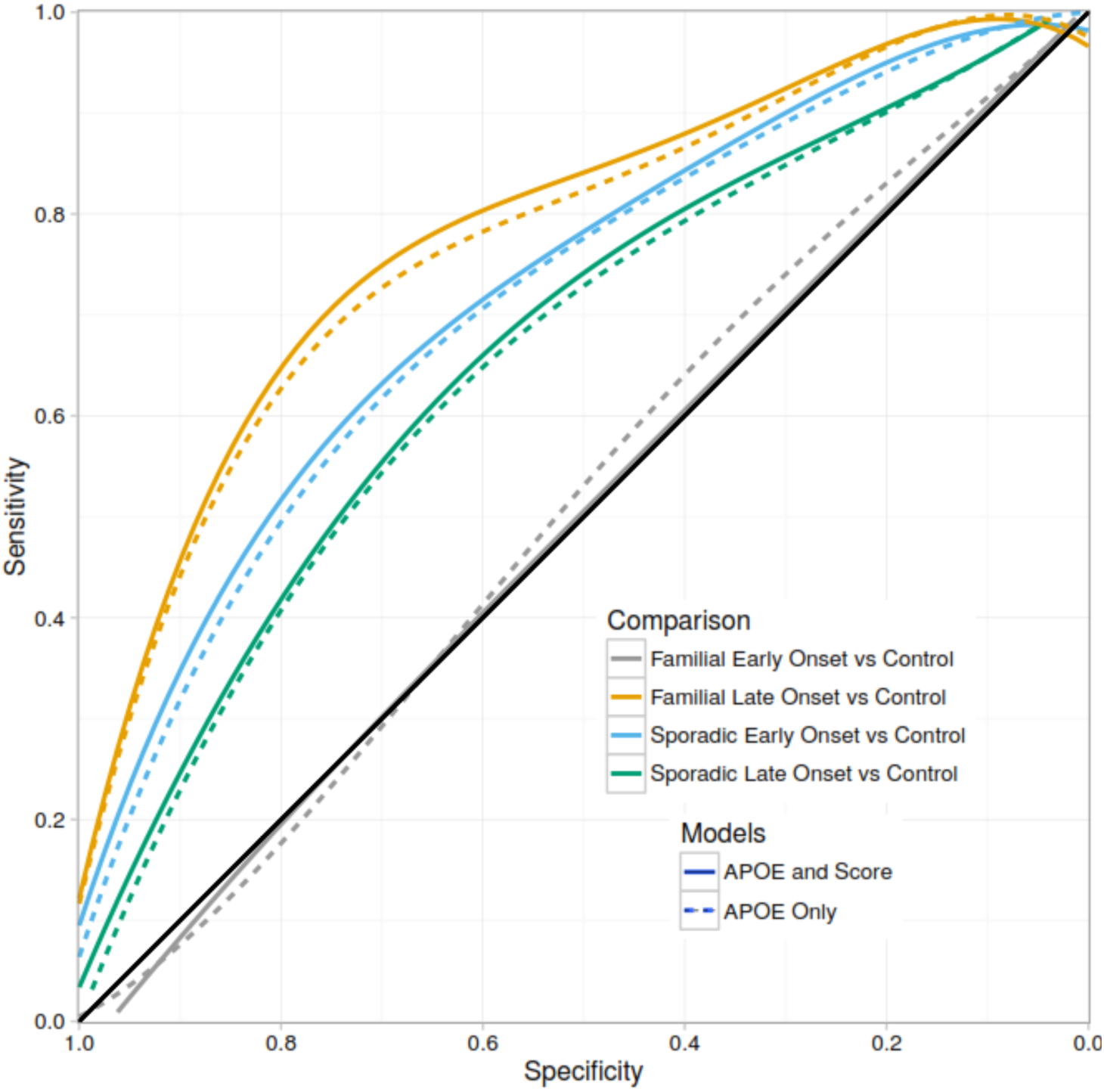
**Receiver operating characteristics – Area under the curve for the different cohorts analyzed.** Dashed lines correspond for the accuracy of obtained for the baseline models that included gender, study, age for late onset and APOE ε4/ε2 alleles. Solid lines reflect the performance of the models that included the effect of the Polygenic Risk Score.

The PRS that summarizes the genetic effect of loci identified for sLOAD, is also significantly associated with risk in the familial late-onset AD (fLOAD) cohort (model 1: OR=1.75; *p*=1.12×10^-7^; Table 2). Indeed, the effect that it has on risk for fLOAD is similar to that seen for *s*LOAD and is not statistically different (*p*=0.13). The model 2 which includes the effects of *APOE* ε2 and ε4 genotypes into the PRS showed an OR=7.85 (p=2.52×10^-48^; **Table 2**) for the fLOAD, which is significantly higher that one calculated for the sLOAD (p=3.01×10^-3^). The AUC for the fLOAD = 0.75 (95% CI=0.73-0,78; **Figure 1**) is significantly higher than the AUC for sLOAD (Venkatraman’s test p<2.2×10-16). This difference is driven by a higher number of *APOE* ε4 carriers in the fLOAD cohort (**Table 1**).

### The effect of the PRS is significantly higher for *s*EOAD than for the *s*LOAD

To study possible effects that the PRS might have for *s*EOAD, we analyzed a cohort of participants with sporadic early onset AD included in the Knight-ADRC and ADNI studies. None of these participants carry a pathogenic mutation in the known AD or FTD genes [7, 41]. We derived the PRS and included elderly non-demented participants as controls. Thus the statistical model corrects for sex, but not age. We observed an increased odds ratio associated with the PRS (model 1: OR=2.27; *p*=1.29×10^-7^; **Table 2**) which was significantly higher than the odds ratio estimated for the *s*LOAD cohort (*p*=9.78×10^-3^). We also observed an increased OR when the effects of *APOE* alleles were included into the PRS (model 2: OR=6.44; p=5.80×10^-26^; **Table 2**). The odds ratio of the PRS with APOE for the sEOAD was also significantly higher than the one calculated for the *s*LOAD (*p*=2.45×10^-2^). Consistently, the ROC analysis revealed a better performance for the *s*EOAD AUC=0.72 (95% CI=0.69-0.76; Figure 1) compared to the *s*LOAD (AUC=0.67, Venkatraman’s test *p*=2.00×10^-3^; **Figure 1**).

In order to confirm that the sEOAD have a higher genetic burden than the sLOAD we compared the PRS of the sEOAD versus the sLOAD directly, instead of comparing the OR. We observed that the PRS for sEOAD is significantly higher than sLOAD (OR=1.47; *p*=9.04×10^-3^; and OR=1.56; *p*=1.95×10^-3^ for the extended model that includes *APOE* effects), suggesting that the known loci could explain in part the earlier onset of these cases. We observed similar results for the analysis of sEOAD cases restricted to an age at onset <60 (data not shown).

### PRS is associated with the age at onset for the sporadic forms of AD

The increased odds ratio that the PRS showed for clinical status when analyzing the *s*EOAD compared to the one obtained for the *s*LOAD led us to question the possible association of PRS with the age at onset (AAO). In particular, we hypothesized that the additive effects of the genome-wide significant loci identified for *s*LOAD would affect the AAO of AD for the entire spectrum of sporadic participants, regardless of the classification of the onset of symptoms (*i*.*e*. early vs late).

We initially verified that in our cohort of *s*LOAD, the PRS is associated with AAO, and as expected a lower AAO is associated with higher PRS for the *s*LOAD (β=-1.20; *p*=7.61×10^-3;^ correcting for cohort and sex and β=-1.66 *p*=7.30×10^-3^ when we included *APOE* effect into the model). In contrast, the analysis within the *s*EOAD cohort did not show a significant association (*p*=0.37; and *p*=0.33 with and without *APOE* effects in the model, respectively).

We also investigated the effect of PRS on AAO in all sporadic AD cases (including all early -and late-onset cases) in specific quantiles of the AAO. We employed quantile regression analysis to model the association of the PRS to 5-quantiles (quintiles) of AAO [42]. This analysis showed that for all of the quintiles the PRS is significantly associated with AAO (p-values are < 0.05; Supplementary Table 1 panel A). Our analyses revealed that all of the coefficient estimates fall within the confidence intervals of the ordinary least squares (OLS) method employed to solve linear regressions (Supplementary Figure 1); indicating: 1) that a linear regression analysis should be sufficient to analyze the association between the PRS and the entire range of AAO (β=-2.10; *p*=1.06×10^-4^); and 2) that this association is not driven exclusively by the cases with older AAO, but all the cases (both early -and late-onset). We observed similar results when *APOE* effects were included into the model (β=-1.99; *p*=1.52×10^-4^; See Supplementary Table 1 panel B for the quantile regression *p-values*; and Supplementary Figure 1).

These analyses indicate that the genetic factors identified for *s*LOAD not only increase the risk for *s*EOAD, but also that their additive effect modulate the entire spectrum of age at onset of sporadic AD participants.

### PRS is not associated with risk for eADAD but shows a significant association with Cerebrospinal fluid (CSF) ptau_181_-Aβ_42_ ratio

We derived the PRS, based on the genetic factors identified for sLOAD, for DIAN participants. The DIAN cohort includes subjects with autosomal dominant AD; and our analyses is restricted to carriers of known pathogenic mutations in *APP, PSEN1* or *PSEN2* genes. We calculated the PRS for these subjects along with elderly non-demented participants, which we employed as controls. In a similar approach to the one we employed for the sEOAD, age was not included in the model, as by design of the test, it predicts perfectly the case-control status. Our analysis did not show any significant association of the PRS with the clinical status for this cohort (model 1: Table 2). Neither did we observe a significant association of the PRS when we incorporated the risk conferred by the *APOE* ε4 and ε2 alleles into the PRS (model 2: Table 2). Nevertheless, we observed that the PRS is associated with the CSF phosphorylated Tau_181_-Aβ_42_ ratio (β=0.18; p=4.36×10^-2^). The CSF ptau_181_-Aβ_42_ ratio was previously shown to be a strong predictor of both the progression of cognitively normal subjects to very mild or mild dementia [43] and the rate of decline across time in individuals with very mild dementia [38]. These results suggest that if any, the effect size of the genetic architecture of *s*LOAD is too small to be significantly identified with statistical power conferred by the cohort of eADAD; but still it also indicates that these genetic risk factors are modulating biological aspects of AD, as it is shown by the association of the PRS and the ptau_181_-Aβ_42_ ratio.

## Discussion

To test for common genetic architecture among autosomal dominant and sporadic forms of early -and late-onset AD, we analyzed four cohorts of well characterized participants; and a common set of elderly non-demented participants. We derived PRS for the participants, assuming an additive non-interaction effect of the common (minor allele frequency (MAF) > 5%) genome-wide significant variants identified for sporadic late onset AD in the IGAP meta-analysis. This approach allowed us to demonstrate a significant association of the additive effect of these genetic factors with the sporadic early-onset and familial late-onset AD. In contrast, we would have required larger samples sizes to perform single SNP association tests and reach levels of significance after correcting for multiple tests. This is because the relatively small effect size of the genetic variants identified to *s*LOAD. The additive effect that additional variants with lower frequency (MAF <5%) associated with AD (i.e. *TREM2* [44, 45], *PLD3* [46], *SORL1* [47, 48] or *ABCA7* [29, 49]) still remains to be analyzed in these cohorts.

Our analysis confirms the hypothesis that the genome-wide significant genetic variants identified for sporadic late onset AD also modulate risk in late-onset familial AD. Furthermore, our analysis showed that the sporadic and familial LOAD participants have similar burden of generic risk factors. However, the samples with a strong family history (fLOAD), independently of the pattern of heritance show the highest percentage of *APOE* ε4 carriers, which is reflected in a higher odd ratio once the effect of this allele is incorporated in the PRS.

Interestingly, the analysis of the sporadic early onset subjects revealed that the PRS, loaded with genetic factors identified from late onset studies, has a more pronounced effect on individuals with younger age at onset. Indeed, this finding not only indicates that the genetic architecture is shared among sporadic cases but also suggests that age at onset of AD is modulated by the additive effect of these loci. We subsequently confirmed this hypothesis identifying a similar effect for different quintiles of the age at onset (Supplementary Table 1; and Supplementary Figure 1).

We could not identify a significant association of the PRS with the early-onset autosomal dominant AD (eADAD) cohort (DIAN). However this cohort has the smallest number of participants. Our empirical power estimates showed that the sample size of this cohort provides a 98% chances to detect an association with odd ratios comparable to that observed for sEOAD. It is more likely that the lack of association of the PRS with eADAD is not because of statistical power, but because the GWAS loci do not confer risk for this population, characterized by the presence of mutations in *APP, PSEN1* and *PSEN2* genes. Nonetheless, we identified an association of the PRS with the CSF ptau_181_-Aβ_42_ ratio; showing that these loci have additional effects on AD pathophysiology most likely in age at onset, but it could also affect disease duration or rate of progression.

The genetic profile summarized in the polygenic risk score should help identify individuals with the highest odds for developing AD dementia, which could benefit the selection of candidates to be present in clinical trials; and also to identify novel therapeutic targets that could modulate the age of symptom onset. Our results indicate that individuals with familial history or earlier-onset are enriched for genetic factors which confer higher odd ratios compared to late-onset cases. These findings can have important repercussion in the design of future genetic studies. Selecting AD cases with earlier onset (but not Mendelian Mutations) or strong family history should provide more statistical power than a similar number of sporadic late-onset cases

One limitation of this study is that to evaluate the extent of overlap of the genetic architectures among the distinct classification of AD we employed common genome-wide significant and replicated variants. A recent study indicates that the SNPs that are significant for AD risk but do not pass the stringent multiple test correction thresholds of GWA studies, can still be informative for the PRS [17]. Additional studies show that low frequency variants, not analyzed in GWA studies are also associated with sLOAD risk [44, 46]. Therefore, further studies including common and rare variants may provide more accurate estimation of the genetic burden of the sEOAD and familial samples in comparison with the sLOAD.

In conclusion, our analysis revealed an overlap among the genetic architecture of the different classifications of AD. The genetic factors identified for sLOAD also affect fLOAD, and sEOAD; with a higher effect size than that observed in the sporadic late-onset form of the disease, and these genetic factors also modulate the age at symptom onset of AD.

## Acknowledgements

We thank all participants and their families for their commitment and dedication to helping advance research into the early detection and causation of AD; and the DIAN research and support staff at each of the participating sites for their contributions to this study.

This work was supported by grants from the National Institutes of Health (R01-AG044546, P01-AG003991, RF1AG053303, R01-AG035083, and R01-NS085419), and the Alzheimer’s Association (NIRG-11-200110). This research was conducted while CC was a recipient of a New Investigator Award in Alzheimer’s disease from the American Federation for Aging Research. CC is a recipient of a BrightFocus Foundation Alzheimer’s Disease Research Grant (A2013359S). The recruitment and clinical characterization of research participants at Washington University were supported by NIH P50 AG05681, P01 AG03991, and P01 AG026276.

Data collection and sharing for this project was supported by The Dominantly Inherited Alzheimer Network (DIAN; U19AG032438, P50 AG-16570 and P50 AG-005142) funded by the National Institute on Aging (NIA) and the German Center for Neurodegenerative Diseases (DZNE). This work was also supported by the NIHR Queen Square Dementia Biomedical Research Unit and the MRC Dementias Platform UK (MR/L023784/1 and MR/009076/1). Exome chip sequencing was supported by the DIAN-TU Pharma Consortium, (the DIAN-TU Pharma Consortium, http://dian-tu.wustl.edu/en/pharma-consortium-members/).

Data collection and sharing for this project was funded by the Alzheimer’s Disease Neuroimaging Initiative (ADNI) (National Institutes of Health Grant U01 AG024904) and DOD ADNI (Department of Defense award number W81XWH-12-2-0012). ADNI is funded by the National Institute on Aging, the National Institute of Biomedical Imaging and Bioengineering, and through generous contributions from the following: AbbVie, Alzheimer’s Association; Alzheimer’s Drug Discovery Foundation; Araclon Biotech; BioClinica, Inc.; Biogen; Bristol-Myers Squibb Company; CereSpir, Inc.; Cogstate; Eisai Inc.; Elan Pharmaceuticals, Inc.; Eli Lilly and Company; EuroImmun; F. Hoffmann-La Roche Ltd and its affiliated company Genentech, Inc.; Fujirebio; GE Healthcare; IXICO Ltd.; Janssen Alzheimer Immunotherapy Research & Development, LLC.; Johnson & Johnson Pharmaceutical Research & Development LLC.; Lumosity; Lundbeck; Merck & Co., Inc.; Meso Scale Diagnostics, LLC.; NeuroRx Research; Neurotrack Technologies; Novartis Pharmaceuticals Corporation; Pfizer Inc.; Piramal Imaging; Servier; Takeda Pharmaceutical Company; and Transition Therapeutics. The Canadian Institutes of Health Research is providing funds to support ADNI clinical sites in Canada. Private sector contributions are facilitated by the Foundation for the National Institutes of Health (www.fnih.org). The grantee organization is the Northern California Institute for Research and Education, and the study is coordinated by the Alzheimer’s Therapeutic Research Institute at the University of Southern California. ADNI data are disseminated by the Laboratory for Neuro Imaging at the University of Southern California.

We thank the Genome Technology Access Center in the Department of Genetics at Washington University School of Medicine for help with genomic analysis. The Center is partially supported by NCI Cancer Center Support Grant #P30 CA91842 to the Siteman Cancer Center and by ICTS/CTSA Grant# UL1TR000448 from the National Center for Research Resources (NCRR), a component of the National Institutes of Health (NIH), and NIH Roadmap for Medical Research. This work was supported by access to equipment made possible by the Hope Center for Neurological Disorders and the Departments of Neurology and Psychiatry at Washington University School of Medicine.

**Scripts and Code:** The R code and scripts we employed to perform the analysis are available in the supplementary information and also in plain text format (file PRS.r)

